# Using publicly available data to conduct rapid assessments of extinction risk

**DOI:** 10.1101/2020.12.14.413906

**Authors:** Michael O. Levin, Jared B. Meek, Brian Boom, Sara M. Kross, Evan A. Eskew

## Abstract

The IUCN Red List plays a key role in setting global conservation priorities. Species are added to the Red List through a rigorous assessment process that, while robust, can be quite time-intensive. Here, we test the rapid preliminary assessment of plant species extinction risk using a single Red List metric: Extent of Occurrence (EOO). To do so, we developed REBA (Rapid EOO-Based Assessment), a workflow that harvests and cleans data from the Global Biodiversity Information Facility (GBIF), calculates each species’ EOO, and assigns Red List categories based on that metric. We validated REBA results against 1,546 North American plant species already on the Red List and found ~90% overlap between REBA’s rapid classifications and those of full IUCN assessments. Our preliminary workflow can be used to quickly evaluate data deficient Red List species or those in need of reassessment, and can prioritize unevaluated species for a full assessment.

## Introduction

The International Union for the Conservation of Nature’s (IUCN; www.iucn.org) Red List is one of the most widely used frameworks to assess extinction risk. The representation of plants on the Red List, however, suffers from pervasive biases that plague conservation science generally (Di Marco et al., 2017; Nic Lughadha et al., 2020). For example, the proportion of described plant species added to the Red List is well below that of vertebrates (10% and 72%, respectively, as of 2020; https://www.iucnredlist.org/resources/summary-statistics, Table 1a). Assessed plants are primarily trees, taxa of particular interest to IUCN Specialist Groups, and species linked to commercial and horticultural interests, among other biases (Bachman et al., 2019; Brummitt, Bachman, & Moat, 2008; Sharrock, 2020). Furthermore, research shows that the Red List may be vastly underestimating plant extinctions, a concerning finding considering that the modern rate of plant extinction is at least 500 times greater than the background extinction rate (Humphreys, Govaerts, Ficinski, Nic Lughadha, & Vorontsova, 2019).

Red listing can also be hampered by features of the assessment process itself. Extant species can be placed into one of six categories: Critically Endangered (CR), Endangered (EN), Vulnerable (VU), Near Threatened (NT), Least Concern (LC), or Data Deficient (DD). The DD category lacks an explicit risk status and is meant to temporarily hold species that lacked sufficient data to be fully assessed. However, already limited conservation resources are rarely diverted to revisit DD listings (Morais et al., 2013), and, as the category has swelled, the estimated cost to fully reassess all DD species is over USD 300 million (Bland et al., 2015). Outside the DD category, many previously assessed species fail to receive mandated regular reassessments; seventeen percent of all assessments were already out of date (> 10 years old) on the 2012 Red List, with the median age of Red List assessments estimated to reach 36 years by 2050 (Rondinini, Di Marco, Visconti, Butchart, & Boitani, 2014).

To address these biases and limitations, several tools have been developed to facilitate rapid, preliminary assessments (Nic Lughadha et al., 2019), particularly for plants (i.e. S. Bachman, Walker, Barrios, Copeland, & Moat, 2020; Callmander, Schatz, & Lowry, 2005; Davis, Govaerts, Bridson, & Stoffelen, 2006; Le Breton et al., 2019; Miller et al., 2013; Utteridge, Nagamasu, Teo, White, & Gasson, 2005). Many of these tools rely upon a single criterion from the full assessment, Criterion B, which focuses on geographic range and is cited in more than 60% of all IUCN assessments (Le Breton et al., 2019). Criterion B relies predominantly upon two measures: Extent of Occurrence (EOO) and Area of Occupancy (AOO). EOO is related to geographic range and measures “the degree to which risks from threatening factors are spread spatially across the taxon’s geographical distribution,” while AOO correlates with population size and approximates a species’ resistance to stochastic events (IUCN Standards and Petitions Committee, 2019; Le Breton et al., 2019). Both have thresholds linked to additional measures of population dynamics and trends that dictate their classification (i.e., if EOO < 100 km^2^, a species could be classified CR).

Previous efforts in this vein have measured the accuracy of an EOO-based assessment method (Miller et al., 2013), used EOO to assess the status of DD plant species (Roberts, Taylor, & Joppa, 2016), and, in one recent instance, produced a streamlined tool that draws from publicly available data sets to identify LC species and submit them to the Red List, allowing the attention of the full assessment to be redirected towards species with higher extinction risk (Bachman et al., 2020). However, few of these rapid assessment frameworks have examined the factors that might influence classification success and none have been tested on large suites of species across broad geographic scales.

Here, we use a publicly available database to gather plant occurrence records for Red Listed species on a continental scale for the first time and analyze the resulting data using a rapid, EOO-based assessment (hereafter, REBA) to assign species a Red List category. We assessed the concordance between our automated classifications and the existing full IUCN classifications, classified DD species into extinction risk categories, and fit statistical models to highlight plant traits and threats that affect the probability of “correct” classification using REBA. Ultimately, our results provide a proof-of-concept for a rapid conservation classification workflow which can be applied to a wide range of species at various scales. This method can serve as a prioritization tool for optimizing resources and effort toward producing full IUCN assessments.

## Methods

### Automated Red List Classification

The REBA workflow begins by using the R package rGBIF (Chamberlain & Boettiger, 2017) to query GBIF for georeferenced occurrence records, which we cap at 50,000 per species to reduce computation time. To further clean the data, we remove records not belonging to kingdom Plantae and filter records to only include “HUMAN_OBSERVATION” or “OBSERVATION” record types to eliminate records that might be georeferenced to a museum location rather than the location of sample collection. We then use the “cc_sea()” function within the R package CoordinateCleaner (Zizka et al., 2019) to remove occurrence records that do not lie over land. Next, REBA uses the R package rCAT to conduct an EOO-based Red List classification (Moat & Bachman, 2020). rCAT calculates EOO as the area of a minimum-convex polygon drawn around known occurrence records (a minimum of 3 is required) and uses IUCN-defined thresholds to classify species as CR, EN, VU, NT, or LC, with EOO values of < 100km^2^, < 5,000km^2^, < 20,000km^2^, < 30,000km^2^, and ≥ 30,000km^2^, respectively. REBA relies exclusively on EOO because there is precedent for such an approach in the literature (see Davis et al., 2006; Miller et al., 2013), and we believe that a metric designed to measure the spatial spread of risk itself (EOO) is more robust for this analysis than one designed to approximate a species’ insurance against that risk (AOO).

### Testing REBA on North American Plant Species

We tested the efficacy of the REBA workflow on each of the 2,662 North American plant species on the Red List. We gathered data on extinction risk, ‘Plant Type’, and ‘Threats’ from the IUCN using the Red List’s advanced search feature (https://www.iucnredlist.org/search; accessed March 23, 2020). After removing 109 species with no GBIF occurrence points and 23 with taxonomic discrepancies, we initially harvested 13,232,845 occurrence records representing 2,530 species. While all of these species are found in North America, not all are native to the continent. Non-native species identified as part of the North American flora by the IUCN were retained for this analysis (hereafter: “North American species”).

After passing through the data cleaning portion of the workflow we were left with 6,566,297 records from 1,829 unique plant species. We joined this occurrence data with Red List assessment data by species and eliminated records from the year of or years following the IUCN’s assessment to ensure REBA was not influenced by data unavailable during the original Red List assessment process. After eliminating species with fewer than 3 cleaned occurrence records, REBA produced EOO-based Red List classifications for 1,546 plant species.

To visualize REBA’s accuracy we generated a tile plot illustrating the overlap between Red List Category classifications generated by the IUCN and by REBA. We then calculated the number of “correctly” classified species (i.e., REBA matched the existing Red List classification), over-classified species (i.e., REBA produced a higher extinction risk category than the existing Red List classification), or under-classified species (i.e., REBA produced a lower extinction risk category than the existing Red List classification). We also utilized the REBA workflow to classify plant species from North America currently classified as DD.

### Statistical Modeling

We derived two smaller data subsets to model the effects of ‘Plant Type’ and ‘Threats’ on the probability of correct classification. The first contained all species that had ‘Plant Type’ data available from the IUCN. In this dataset, we condensed the 18 original ‘Plant Type’ categories assigned by the IUCN into ten new biologically relevant categories (Appendix 1), effectively eliminating those that would otherwise be represented by a limited sample of species. This modeling dataset contained 1,533 plant species. The second dataset represented all species that had ‘Threats’ data available from the IUCN. ‘Threats’ categories remained unaltered, and the dataset contained 396 species.

We then constructed two separate multilevel Bayesian models with a binomial outcome distribution where correct classification was the variable of interest. Both models included the number of occurrence points as a main effect. One included ‘Plant Type’ categories as varying effects, and the other included ‘Threats’ categories as varying effects. These models estimate the effect of number of occurrence points, ‘Plant Type’, and ‘Threats’ on REBA’s probability of correct classification. We specified and fit our models using the “map2stan()” function within the ‘rethinking’ R package (Carpenter et al., 2017; McElreath, 2016). We fit each model using four independent MCMC chains, specifying 7,500 total model iterations, 2,500 of which were considered warmup. As a result, our model inferences are based on 20,000 posterior samples from each model (5,000 post-warmup samples per chain with four chains total). After model fitting, we inspected parameter trace plots and R-hat values to confirm convergence and good model fits (Gelman & Rubin, 1992). We report parameter estimates using posterior means and 99% highest posterior density intervals (HPDIs) where appropriate.

### Counterfactual Predictions of Classification Accuracy

We visualized our model results with counterfactual plots using the full posterior probability distributions to show the implied relationships between the probability of correct classification and ‘Plant Type’ or ‘Threats’, respectively. We plotted the implied probability of correct classification using four occurrence point sample sizes: 100, 1,000, 10,000, and 20,000. In effect, we imagine applying REBA for a species with a given ‘Plant Type’ or ‘Threats’ classification across a range of arbitrary sample sizes, where the implied probability of correct classification is informed by the observed data that was used to fit the model.

The inference derived from our fit models can also be applied to DD species, allowing us to generate posterior distributions for the implied probability of correct Red List classification analogous to the counterfactual analyses just described. All 13 DD species with REBA-generated classifications had ‘Plant Type’ data, but only 5 had ‘Threats’ data. Therefore, we used the full posterior distributions from our ‘Plant Type’ model to generate the implied probability of correct classification for each of these species, given their ‘Plant Type’ and the actual number of cleaned occurrence points available. Thus, our analyses for DD species represent the implied probability of correct classification for the exact species-level data as analyzed through REBA.

## Results

### Classification Overlap and Modeling Probability of Correct Classification

REBA correctly classified 1,379 of 1,533 species (89.95%) in our dataset of North American plant species. An overwhelming majority of correct classifications (99.49%) were for LC species (Fig. 1). We under-classified 58 species (3.78%) and over-classified 96 (6.26%).

**Fig. 1.**
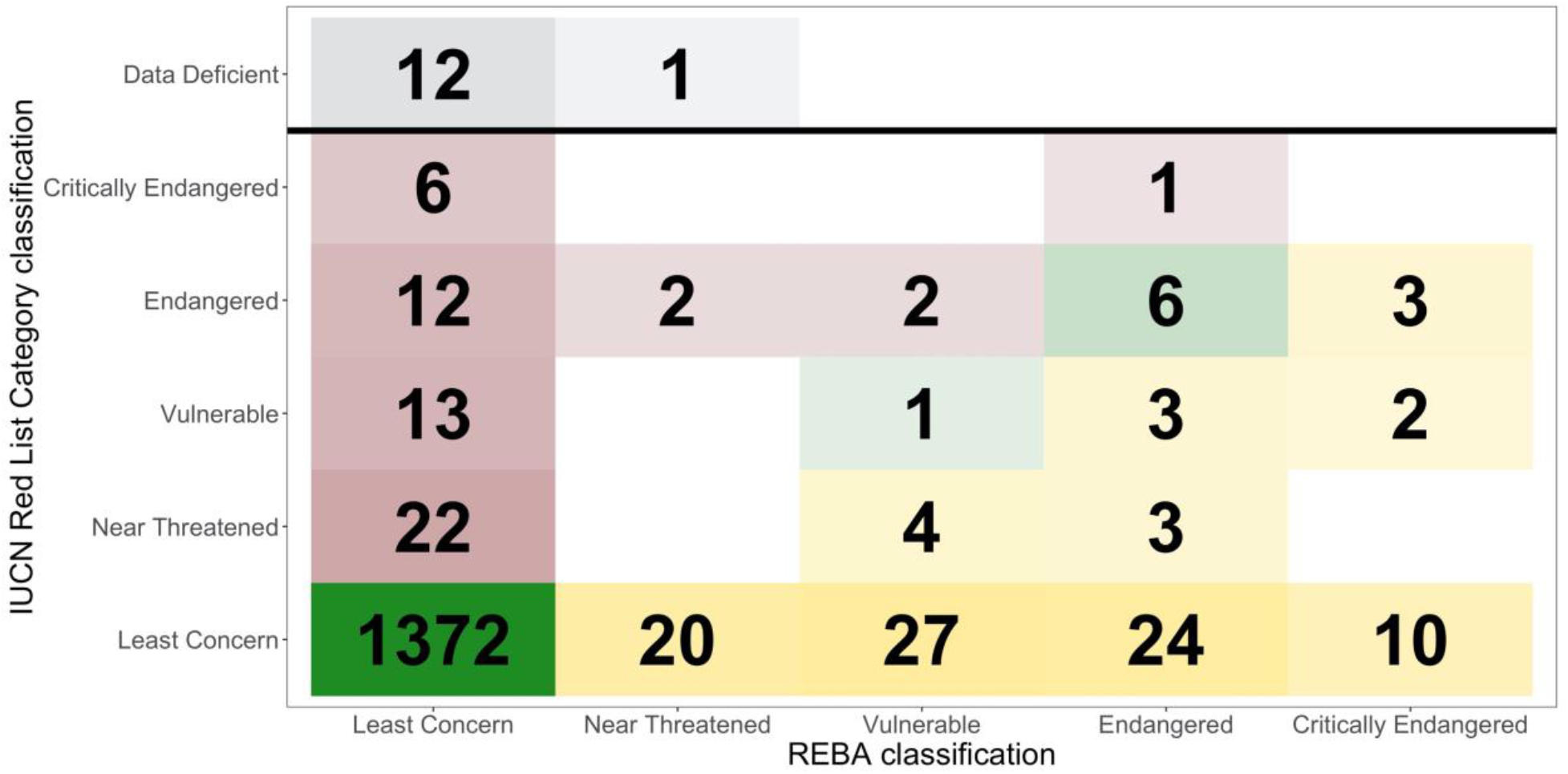
Below the bold line, each tile represents an intersection of IUCN threat category classifications: those assigned by the IUCN and those assigned using REBA. Green tiles along the diagonal represent matching classifications, where both the IUCN and REBA classified species into the same categories. Yellow tiles below the diagonal are those we over-classified, where REBA placed species into a higher extinction risk category than that produced by the IUCN’s classification. Red tiles above the diagonal represent those species we under-classified, where REBA placed species into a lower extinction risk category than that produced by the IUCN’s classification. Tile transparency is a function of the number of species associated with that classification combination. Grey tiles above the bold line represent the threat categories into which we classified 13 DD species using REBA.

‘Geophytes’ contained the highest proportion of under-classified species among ‘Plant Type’ (17.02%; Fig. 2) and exhibited the strongest negative effect on the probability of correct classification (mean effect on logit scale [99% HPDI]: −1.14 [−2.25, 0.00]; Fig. 3). ‘Annuals’, ‘Ferns’, and ‘Graminoids’ contained no under-classified species (0%; Table S1, Fig. 2) and ‘Annuals’ exhibited the strongest positive effect (1.13 [−0.22, 3.48]; Fig. 3). In the ‘Plant Type’ model, posterior estimates for effect of the number of points (NOP) maintained support for positive values across the entire 99% HPDI (1.60 [0.63, 3.06]).

**Fig. 2.**
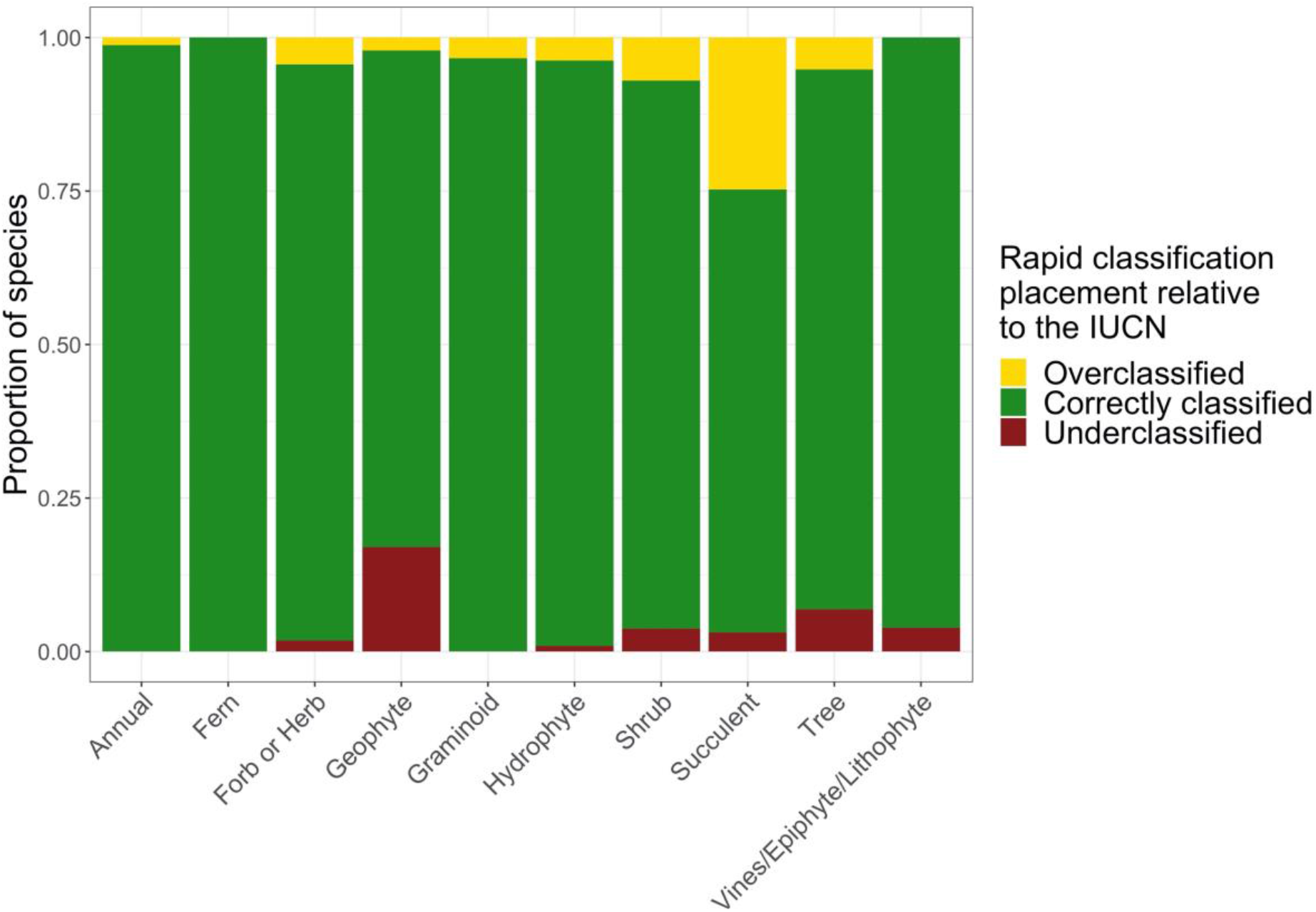
Each bar represents the total number of species assigned to a given ‘Plant Type’ by the IUCN. The proportion of those species that were classified correctly, over-classified, and under-classified are represented respectively by green, yellow, and red sections. It is important to know that some species are counted across multiple plant types, as they were assigned more than one by the IUCN (i.e., *Acer grandidentatum* is classified as both a ‘Shrub’ and ‘Small Tree’). See Table S1 for raw data.

**Fig. 3.**
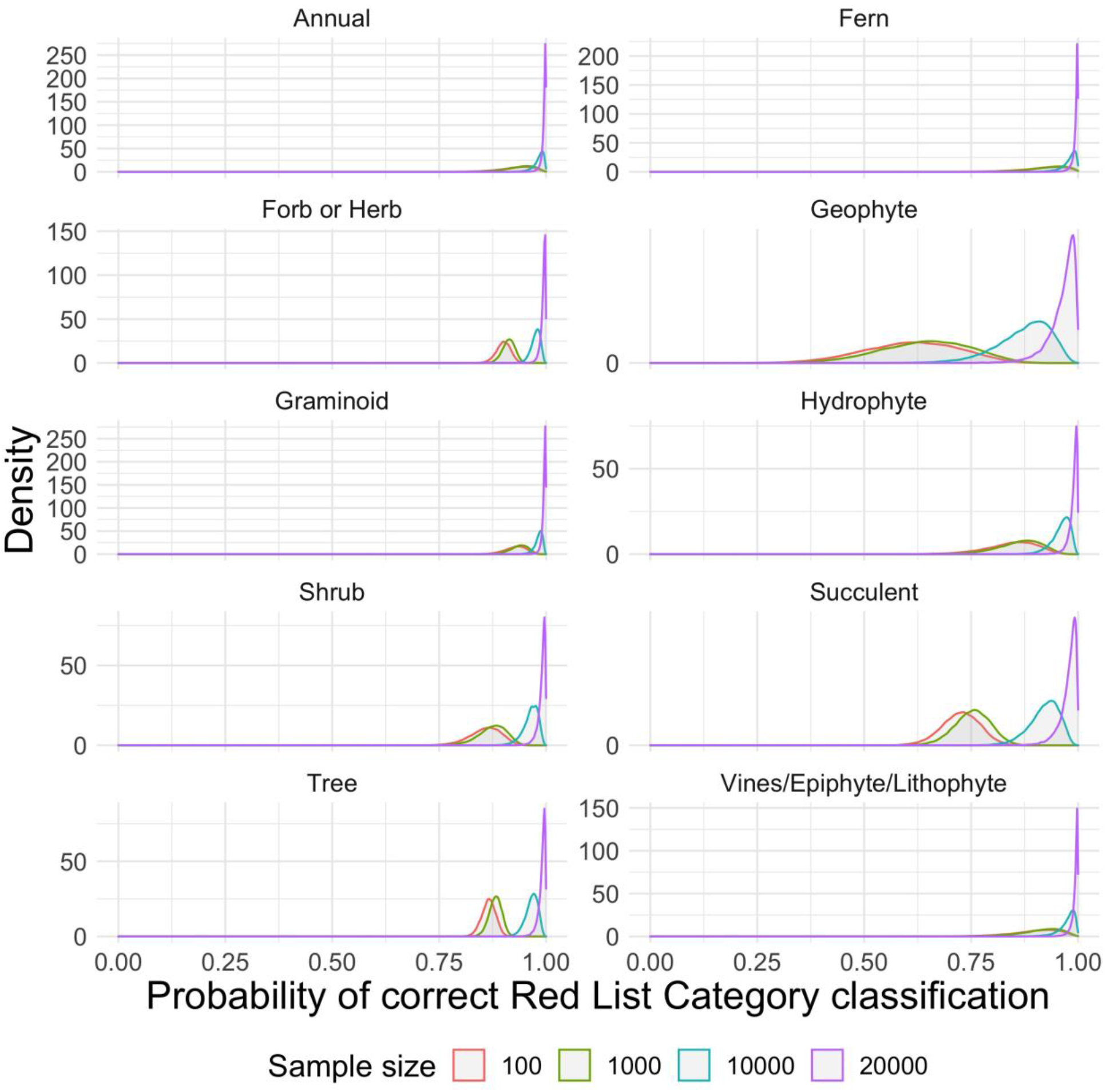
Posterior probability distributions representing the influence of IUCN assigned ‘Plant Type’ on the probability of correct REBA classification at different sample sizes.

Among ‘Threats’ categories, ‘Human Intrusions and Disturbance’ had the highest proportion of under-classified species (27.03%; Fig. 4), while ‘Residential and Commercial Development’ exhibited the most negative effect on the probability of correct classification (−0.55 [−1.20, 0.03]; Fig. 5). ‘Climate Change and Severe Weather’ had the lowest proportion of under-classified species (12.62%; Table S2, Fig. 4) and exhibited the largest positive effect on correct classification (0.24 [−0.33, 0.91]; Fig. 5). In the ‘Threat’ model, posterior estimates for NOP again had support constrained to only positive values (0.99 [0.16, 2.26]).

**Fig. 4.**
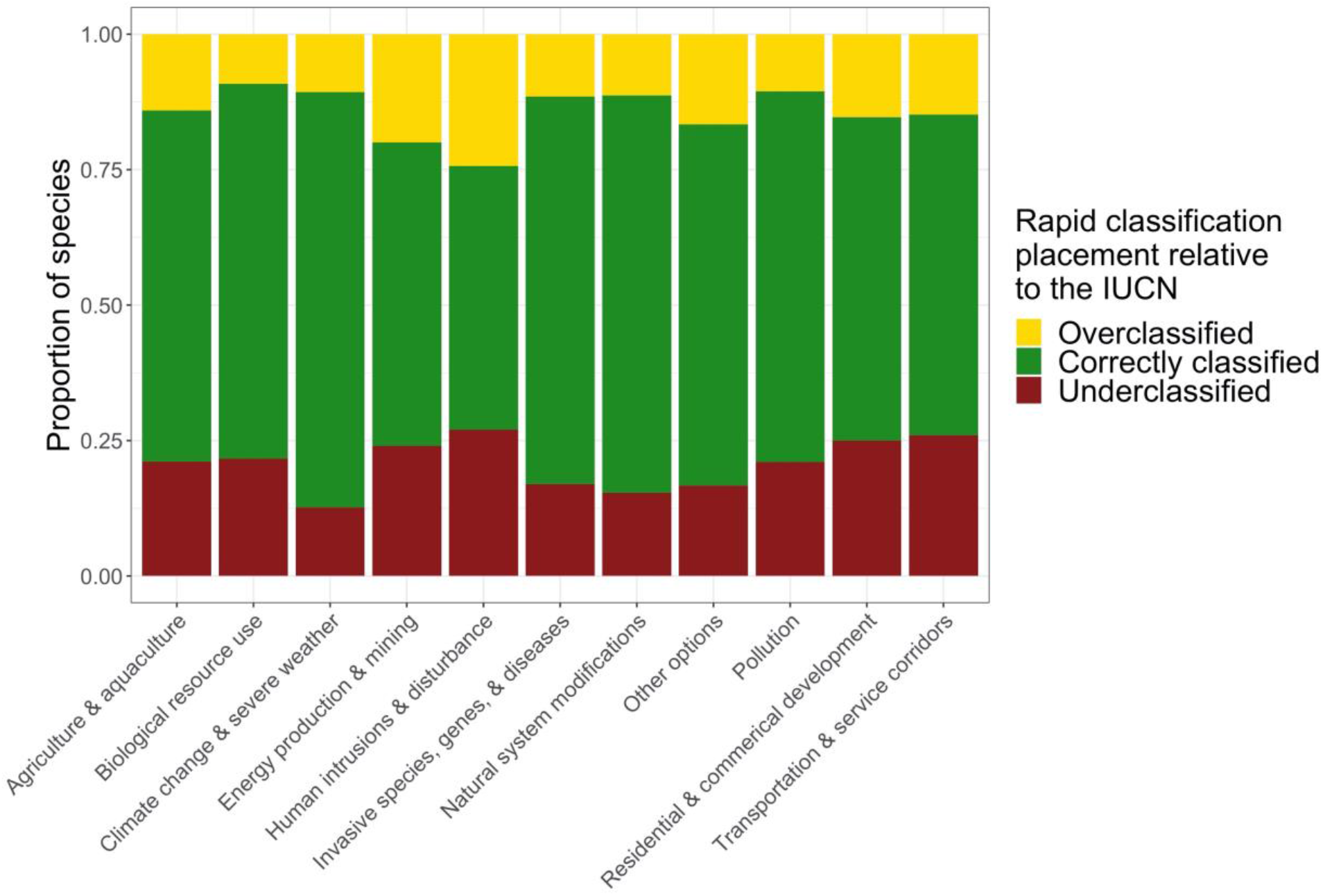
Each bar represents the total number of species assigned to a given ‘Threats’ category by the IUCN. The proportion of those species that were classified correctly, over-classified, and under-classified are represented respectively by green, yellow, and red sections. It is important to know that some species are counted across multiple threat types, as they were assigned more than one by the IUCN (i.e., *Acer rubrum* is classified as threatened by ‘Natural System Modifications,’ ‘Invasive and Other Problematic Species, Genes, and Disease,’ and ‘Climate Change and Severe Weather’). See Table S2 for raw data.

**Fig. 5.**
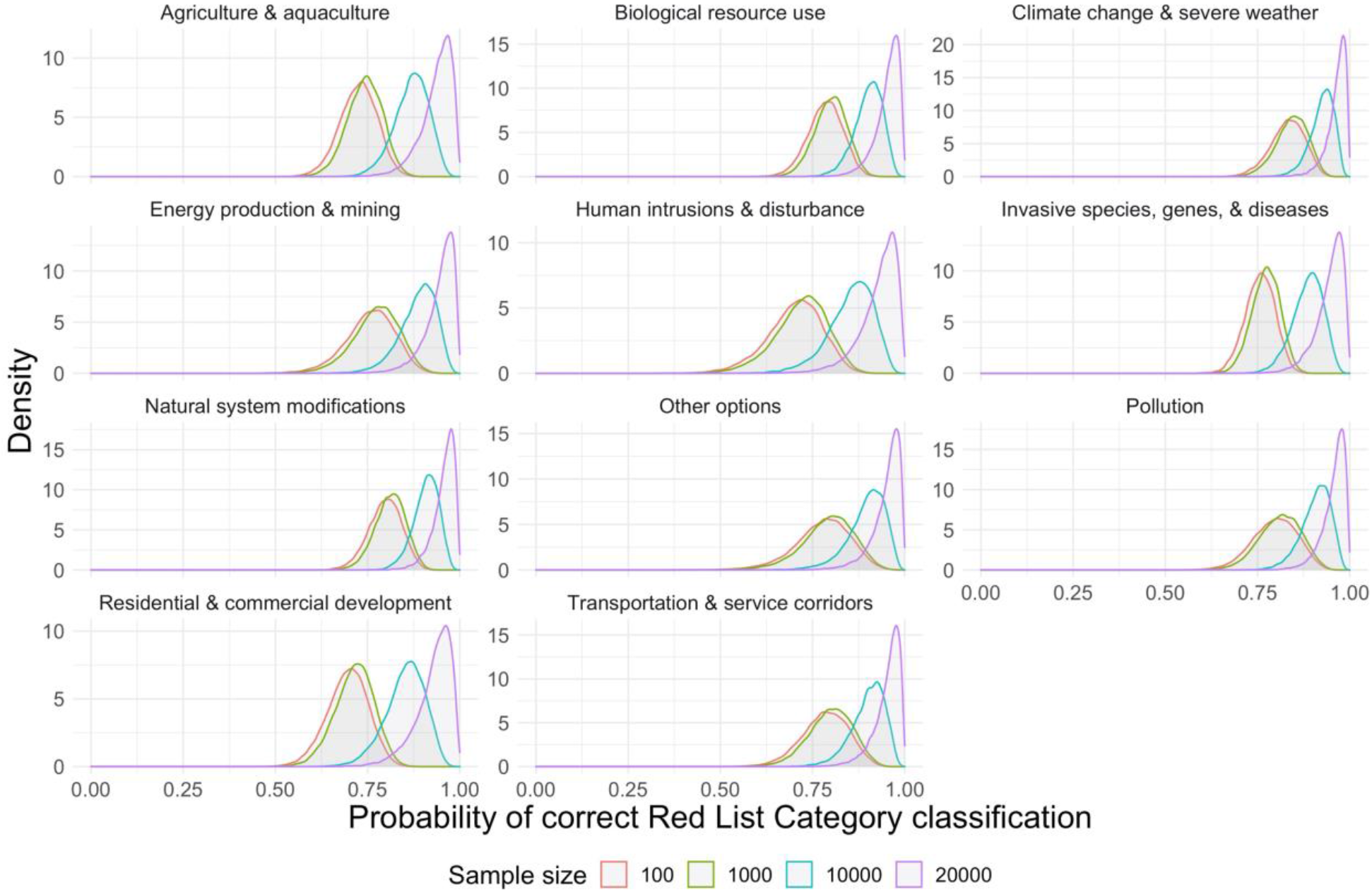
Posterior probability distributions representing the influence of IUCN assigned ‘Threats’ on the probability of correct REBA classification at different sample sizes.

### Classifying Data Deficient Species

Thirteen North American species classified by the IUCN as DD remained after data filtering (Table S3). REBA classified twelve as LC and one as NT. Their mean EOO was 56,612,597.68 km^2^, with a minimum of 22,002.41 km^2^ and a maximum of 162,676,407.50 km^2^. Their mean NOP was 4,322.80, with a minimum of 9 and a maximum of 40,298. All DD species had mean implied probabilities of correct classification above 0.70 (Table S3, Fig. 6). *Ulmus glabra* had the highest mean implied probability of correct classification (1.00 [.99, 1.00]) and *Zingiber zerumbet* had the lowest (.73 [.49, .90]; Table S3, Fig. 6). These thirteen species represent a third of the total number of North American species classified as DD (n = 39). The 26 species that were removed in our data cleaning process contained fewer than three cleaned occurrence records, precluding an EOO calculation. They remain priorities for further data collection.

**Fig. 6.**
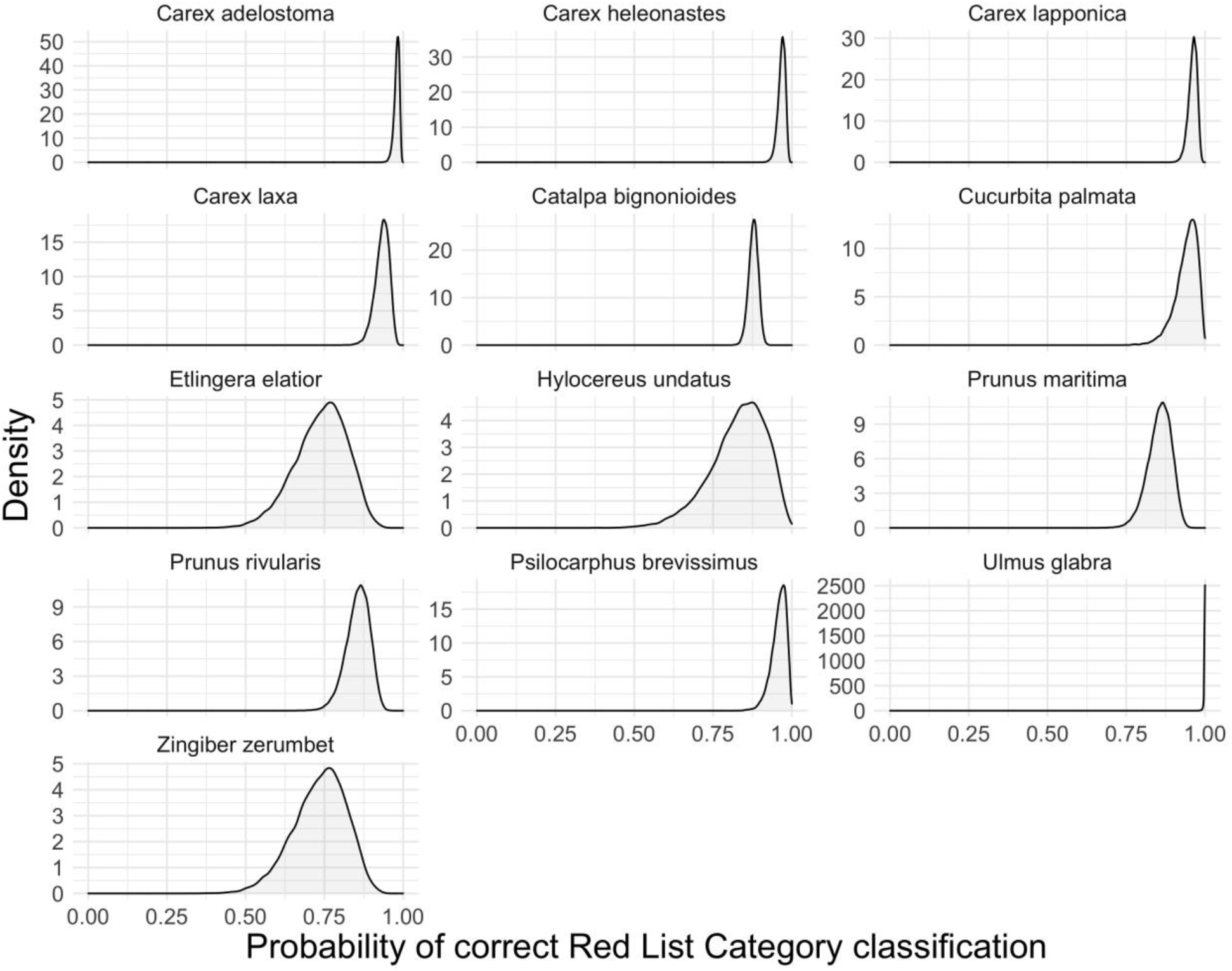
Posterior probability distributions representing the probability of correct REBA classification of 13 DD species. We employ the full posterior distributions of the ‘Plant Type’ model to produce the implied probability of correct classification for each of these species based on their ‘Plant Type’ and the actual number of cleaned occurrence points available for that species. See Table S3 for raw data.

## Discussion

REBA can swiftly and accurately produce preliminary Red List assessments on a continental scale that match existing Red List assessments approximately 90% of the time. Our modeling efforts indicated that the number of points available for a species is the most important contributor to the probability of correct classification. REBA classified 13 DD species into non-threatened categories with more than 70% probability of presumed correct classification. Results from the ‘Threats’ model are less easily interpreted than those of the ‘Plant Type’ model. The mean number of ‘Plant Type’ per species was 1.29, while for ‘Threats’ it was 2.35. Because many species are affected by multiple threats, it is more difficult to parse individual effects of each one.

One of our most significant concerns regarding REBA came from under-classified species, which represent species of current conservation concern that would be masked under this assessment framework. REBA under-classified all seven North American species listed as CR, labelling six as LC: five *Fraxinus* species threatened by the Emerald Ash Borer (*Agrilus planipennis*), and the American Chestnut (*Castanea dentata*), threatened by the chestnut blight caused by *Cryphonectria parasitica*. These are widespread species, and their large EOOs mask the tremendous risk that invasive species and disease represent across their range. These particular threats are thus potential confounders of the REBA framework, particularly for species with large EOOs. This matches observations made in other applications of similar methods (S. Bachman et al., 2020).

Another concern arises from the data itself. We found that the probability of correct classification increases substantially across all ‘Plant Types’ and ‘Threats’ with increasing NOP: REBA assessments based on > 20,000 records approach a 100% probability of correct classification (Fig. 3, Fig. 5). However, including more data may expose REBA to well-founded data quality concerns for publicly available data (see Meyer, Weigelt, & Kreft, 2016). Our data cleaning relied primarily on GBIF metadata and only implemented spatial filters to remove non-terrestrial points. Bachman et al. used a similar methodology but employed a stricter spatial filter that limited occurrence data to those records that overlapped with a species’ native range (Bachman et al., 2020). However, such strict filtering potentially masks valuable occurrence records of non-native but naturalized species. Critical spatial filtering and further assessment of more rigorous cleaning methods to improve data quality is central to further refinement of rapid assessment tools.

While acknowledging these concerns, REBA did produce correct classifications in ~90% of cases. The majority were for LC species, which could be a result of geographic bias in our study; North American species are well-studied and likely to have significant data available (Meyer et al., 2016), which may underlie the high number of LC classifications that REBA produced. By a similar logic, the more occurrence points a species has, the more likely it is to be of minimal conservation concern–a species with > 10,000 records is likely to be widespread. However, the identification of LC species is valuable for assessment prioritization. Once recognized, LC species can be put aside in the assessment pipeline in favor of those more likely to have a higher extinction risk that would benefit from more immediate attention (Bachman et al., 2020).

REBA’s optimal function is to work alongside the full Red List assessment process, helping to focus limited human and financial capital on species in need of immediate attention by discovering species of least concern. We have demonstrated that it can accurately and rapidly leverage publicly available data to operate on a continental scale to prioritize plant conservation efforts (Antonelli et al., 2020). It can be applied to swiftly narrow the growing pool of DD species, address the growing backlog of species in need of reassessment, and provide a preliminary pass for unassessed species. Further refinement of REBA, as well as broader spatial and taxonomic applications, are necessary and welcomed. The need for action is immediate–there is little time to waste.

## Data Accessibility Statement

Data downloaded from the IUCN, along with R scripts used for GBIF data collection, filtering, analyses, and visualization are freely available on GitHub: https://github.com/eveskew/plant_rapid_assessment.

## Appendix 1: Rationale for grouping of ‘Plant Types’

We modeled the probability of correct classification among different ‘Plant Types’, which were previously defined by the IUCN (https://www.iucnredlist.org/resources/classification-schemes; Plant and Fungal Growth Forms Classification Scheme). The 1,546 species used to validate the REBA classification fell into 18 of the 24 IUCN ‘Plant Type’ categories. We combined some categories to improve statistical analysis and visualization, and these combinations are justified both biologically and by IUCN’s ‘Plant Type’ classification guidelines. For example, IUCN has multiple ‘Tree’ categories differentiated by size (i.e. ‘Tree - size unknown’, ‘Tree - large’, ‘Tree - small’), but admits that the categories based on size are “sub-types which may be dropped at some point in the future”, so we combined these categories into one ‘Tree’ category. Sub-categories for both ’Succulent’ and ‘Shrub’ categories were combined for similar reasons. We combined the ‘Vine’, ‘Epiphyte’, and ’Lithophyte’ categories due to low sample sizes, which we thought was biologically reasonable because plants of these types often grow on atypical substrates and tend to display a climbing habit.

