## Supplementary material for "Using publicly available data to conduct rapid assessments of extinction risk": Table S1

**Table S1.** Data on the ‘Plant Type’ associated with a given species was obtained from the IUCN alongside species assessment data. The under-classified species were of special concern, as their true extinction risk would be masked by REBA. A rationale for the condensation of ‘Plant Type’ categories can be found in Appendix 1. Note that many species had several associated plant types and were counted multiple times as correctly, over-, or under- classified within each plant type they were assigned.

| | Over-classified | | Correctly Classified | | Under-classified | |
|---|---|---|---|---|---|---|
| Plant Type | Number of Species | Proportion | Number of Species | Proportion | Number of Species | Proportion |
| Annual | 1 | 1.22% | 81 | 98.78% | 0 | 0.00% |
| Fern | 0 | 0.00% | 24 | 100.00% | 0 | 0.00% |
| Graminoid | 8 | 3.39% | 228 | 96.61% | 0 | 0.00% |
| Hydrophyte | 4 | 3.74% | 102 | 95.33% | 1 | 0.93% |
| Forb or Herb | 27 | 4.39% | 577 | 93.82% | 11 | 1.79% |
| Succulent | 24 | 24.74% | 70 | 72.16% | 3 | 3.09% |
| Shrub | 15 | 7.04% | 190 | 89.20% | 8 | 3.76% |
| Vines/Epiphyte/Lithophyte | 0 | 0.00% | 25 | 96.15% | 1 | 3.85% |
| Tree | 30 | 5.17% | 510 | 87.93% | 40 | 6.90% |
| Geophyte | 1 | 2.13% | 38 | 80.85% | 8 | 17.02% |
