## Supplementary material for "Using publicly available data to conduct rapid assessments of extinction risk": Table S2

**Table S2.** Data on the ‘Threats’ associated with a given species was obtained from the IUCN alongside species assessment data. The under-classified species were of special concern, as their true extinction risk would be masked by REBA. Note that many species had several associated threats and were counted multiple times as correctly, over-, or under- classified within each threat they were assigned.

| | Over-classified | | Correctly Classified | | Under-classified | |
|---|---|---|---|---|---|---|
| Threat | Number of Species | Proportion | Number of Species | Proportion | Number of Species | Proportion |
| Climate change & severe weather | 11 | 10.68% | 79 | 76.70% | 13 | 12.62% |
| Natural system modifications | 17 | 11.33% | 110 | 73.33% | 23 | 15.33% |
| Other options | 1 | 16.67% | 4 | 66.67% | 1 | 16.67% |
| Invasive species, genes, & diseases | 19 | 11.52% | 118 | 71.52% | 28 | 16.97% |
| Pollution | 4 | 10.53% | 26 | 68.42% | 8 | 21.05% |
| Agriculture & aquaculture | 20 | 14.08% | 92 | 64.79% | 30 | 21.13% |
| Biological resource use | 11 | 9.17% | 83 | 69.17% | 26 | 21.67% |
| Energy production & mining | 5 | 20.00% | 14 | 56.00% | 6 | 24.00% |
| Residential & commercial development | 19 | 15.32% | 74 | 59.68% | 31 | 25.00% |
| Transportation & service corridors | 4 | 14.81% | 16 | 59.26% | 7 | 25.93% |
| Human intrusions & disturbance | 9 | 24.32% | 18 | 48.65% | 10 | 27.03% |
