## Supplementary material for "Using publicly available data to conduct rapid assessments of extinction risk": Table S3

**Table S3.** Proposed classifications for the 13 North American Data Deficient species that survived our data cleaning procedure and mean posterior distributions for the implied probability of correct Red List Category classification with 99% HPDI.

| | Classification Results and Summary Information | | | | Modeling Results | | | |
|---|---|---|---|---|---|---|---|---|
| **Data Deficient Species** | **IUCN Assessment Year** | **EOOcat** | **EOOkm2** | **NOP** | **Mean** | **Standard Deviation** | **0.50%** | **99.50%** |
| *Ulmus glabra* | 2017 | LC | 107,307,670.00 | 40298 | 1.00 | 0.00 | 0.99 | 1.00 |
| *Carex adelostoma* | 2015 | LC | 871,098.93 | 4946 | 0.98 | 0.01 | 0.95 | 0.99 |
| *Carex heleonastes* | 2015 | LC | 21,711,348.30 | 1782 | 0.97 | 0.01 | 0.93 | 0.99 |
| *Carex lapponica* | 2015 | LC | 9,159,336.21 | 549 | 0.96 | 0.01 | 0.92 | 0.99 |
| *Psilocarphus brevissimus* | 2015 | LC | 509,453.97 | 9 | 0.96 | 0.02 | 0.88 | 1.00 |
| *Cucurbita palmata* | 2017 | LC | 136,197.98 | 51 | 0.94 | 0.04 | 0.81 | 1.00 |
| *Carex laxa* | 2015 | LC | 10,173,930.50 | 688 | 0.93 | 0.02 | 0.86 | 0.98 |
| *Hylocereus undatus* | 2009 | LC | 162,676,408.00 | 74 | 0.83 | 0.09 | 0.54 | 0.98 |
| *Prunus rivularis* | 2014 | LC | 1,534,379.61 | 11 | 0.86 | 0.04 | 0.74 | 0.93 |
| *Prunus maritima* | 2014 | NT | 22,002.41 | 9 | 0.86 | 0.04 | 0.74 | 0.93 |
| *Catalpa bignonioides* | 2018 | LC | 147,874,052.00 | 801 | 0.88 | 0.02 | 0.84 | 0.91 |
| *Etlingera elatior* | 2018 | LC | 157,333,989.00 | 163 | 0.74 | 0.08 | 0.50 | 0.91 |
| *Zingiber zerumbet* | 2018 | LC | 116,653,903.00 | 30 | 0.73 | 0.08 | 0.49 | 0.90 |
